# A cohort-based global sensitivity benchmark of MRI-derived whole-heart electromechanical models in healthy hearts

**DOI:** 10.64898/2026.03.27.714701

**Authors:** Shahrokh Rahmani, Jim Pouliopoulos, Angela W. C. Lee, Rosie K. Barrows, Jose Alonso Solis-Lemus, Marina Strocchi, Cristobal Rodero, Abdul Qayyum, S. Samaneh Lashkarinia, Caroline Roney, Christoph M. Augustin, Gernot Plank, Diane Fatkin, Andrew Jabbour, Steven A. Niederer

## Abstract

Patient-specific four-chamber electromechanical models provide a physics-constrained framework for investigating whole-heart cardiac physiology and disease mechanisms. Identifying which model parameters impact whole-heart function is important for understanding cellular-, tissue-, and organ-scale determinants of cardiac performance and for calibrating patient-specific models. However, previous global sensitivity analyses of cardiac electromechanical models have typically been performed on a single heart, and systematic evaluation of how parameter influence compares across anatomically different subjects remains limited.

We created four-chamber electromechanical models using cardiac MRI from five healthy subjects (*n* = 5). The models simulated atrial and ventricular cellular electrophysiology, calcium dynamics, and active contraction, with heterogeneous fibre orientation, transversely isotropic tissue mechanics, pericardial constraint, and a closed-loop cardiovascular system providing physiological boundary conditions. In total, 46 parameters described the integrated model.

Using Gaussian process emulators, we performed multi-scale global sensitivity analysis to evaluate the relative contribution of model parameters to left and right atrial and ventricular function. Across all anatomies, the most influential parameters were systemic and pulmonary resistances, ventricular end-diastolic pressures, and the venous reference pressure, highlighting the dominant role of haemodynamic loading conditions in governing pressure- and volume-based outputs.

A chamber-level analysis of atrioventricular coupling revealed a phase-dependent pattern. Atrial pressures were predominantly governed by global haemodynamic parameters (*>* 90% of total sensitivity), atrial filling volumes showed substantial ventricular influence (≈40–55% across anatomies), and atrial end-systolic volumes were primarily determined by intrinsic atrial parameters (≈60–65%). These patterns were consistent across subjects despite differences in anatomy.

We show that, in healthy male subjects, inter-individual anatomical variation does not substantially change the ranking of dominant parameters. This work provides a repeatable modelling and sensitivity analysis framework and establishes a benchmark reference for whole-heart electromechanical modelling in healthy hearts.

**Author summary:** Computational models of the heart can simulate cardiac physiology in unprecedented detail, but these models contain many parameters whose influence on predicted function is not fully understood. We built patient-specific four-chamber heart models from MRI scans of five healthy subjects and used statistical methods to systematically test how 46 model parameters affect simulated cardiac performance. Across all five subjects, we found that the haemodynamic loading parameters, including systemic and pulmonary vascular resistance, ventricular filling pressures, and the venous reference pressure, consistently had the greatest influence on the model outputs, regardless of differences in individual heart anatomy. This finding suggests that in healthy resting conditions, the boundary conditions of the cardiovascular system, rather than individual differences in heart geometry or electrical properties, are the primary drivers of whole-heart function. We also found a structured coupling pattern between the upper and lower heart chambers, where global haemodynamic parameters dominate atrial pressure regulation, ventricular mechanics shape atrial filling, and intrinsic atrial properties control atrial emptying. This work provides a benchmark dataset of five anatomically detailed heart models and a sensitivity analysis framework to guide calibration of future cardiac digital twin models.

## Introduction

Computational cardiac models are increasingly used for structural and functional analysis of the cardiovascular system [1]. Beyond descriptive characterisation, these models enable mechanistic investigation of disease processes, support patient stratification and outcome prediction, and provide simulation-based evaluation and optimisation of interventions [2–5].

Recent patient-specific cardiac modelling studies have focused on establishing robust workflows for constructing anatomically detailed computational heart models directly from clinical imaging data, including segmentation, mesh generation, and personalisation of model geometry [6–8]. These image-based models represent patient-specific digital twins of the heart, integrating anatomical structure with biophysical simulation to reproduce individual cardiac physiology. These studies demonstrated the feasibility and reliability of image-based heart model reconstruction across a range of cardiac conditions, providing a foundation for the integration of clinical imaging with physics-based cardiac simulation [6, 9]. More recent studies have extended these approaches to clinically meaningful patient cohorts, demonstrating the predictive power of digital twins for patient-specific mechanisms and therapy guidance, particularly in arrhythmia and ablation planning [10]. Large-scale efforts have emerged to enable the generation of cardiac digital twins across large populations [11]. Nevertheless, despite these advances, systematic evaluation of how model behaviour and parameter influence compare across multiple anatomically different subjects, remains relatively limited. Addressing this gap is essential for understanding the consistency of findings from the models across individuals and for supporting interpretation at the cohort level.

The impact of individual parameters on whole-heart function can be investigated using sensitivity analysis. Global sensitivity analysis (GSA) explores the influence of parameters across the full parameter space and accounts for interactions between parameters, in contrast to local sensitivity approaches that evaluate behaviour around a single parameter set. In cardiac digital twin models, which often contain a large number of parameters, GSA is particularly valuable for identifying the most influential parameters and guiding patient-specific model calibration. However, variance-based GSA methods typically require a large number of simulations, making direct application to computationally expensive electromechanical heart models challenging. Surrogate modelling approaches, such as Gaussian process emulators (GPEs), provide an efficient strategy to overcome this limitation. A study in a heart failure patient model identified influential parameters [12], while more recent works have extended GSA to four-chamber electromechanics [13], haemodynamics using surrogate models [14, 15], and uncertainty quantification in electrophysiology [15]. However, these studies all performed their sensitivity analysis on a single heart, so it is not clear if these results generalise across individuals with different cardiac anatomies.

In this study, we apply an established image-based modelling workflow to construct four-chamber whole-heart electromechanical models from patient-specific anatomy derived from cardiac MRI. We apply GSA across a cohort of five healthy subjects to address three related questions: first, whether the influence of key parameters on model outputs remains consistent across different hearts despite subject-specific variations in cardiac geometry; second, whether the dominant parameter groups identified in a heart failure model [13] are preserved or altered in healthy anatomies; and third, given the strong physiological coupling between atria and ventricles, how atrial function in healthy hearts decomposes into ventricular-specific, atrial-specific, and global (haemodynamic) parameter contributions when considering all chamber-related inputs across electrophysiological, mechanical, and calcium-handling subsystems, thereby providing a comprehensive assessment of atrioventricular coupling.

## Materials and methods

### Cohort

Five healthy subjects, recruited as part of an alcohol consumption clinical study, were selected from Cine Magnetic Resonance Imaging (MRI) datasets acquired at Victor Chang Cardiac Research Institute (VCCRI). Patient recruitment and data acquisition were conducted at St Vincent’s Hospital, Sydney, and were approved by the St Vincent’s Hospital Human Research Ethics Committee, Sydney, New South Wales, Australia (2021/ETH00654). All patients provided written informed consent. Demographic details of the subjects are provided in Table 1.

**Table 1.**
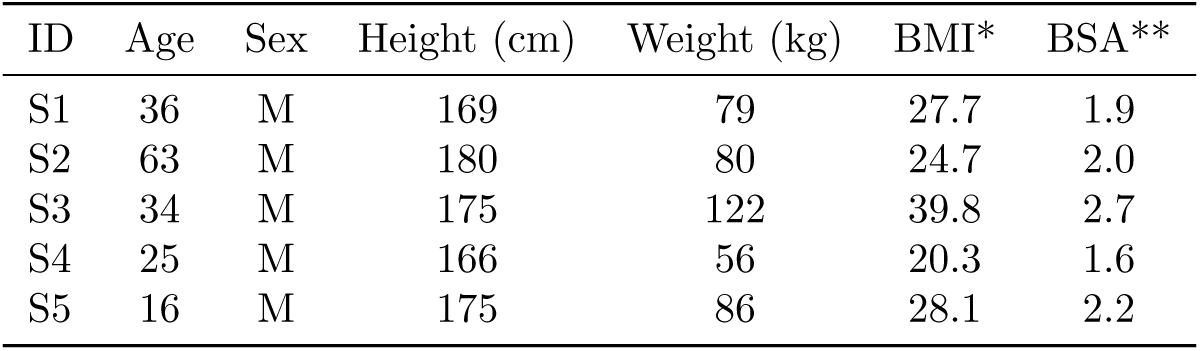
Demographic details of the five subjects; *BMI: Body Mass Index, **BSA: Body Surface Area

| ID | Age | Sex | Height (cm) | Weight (kg) | BMI* | BSA** |
| --- | --- | --- | --- | --- | --- | --- |
| S1 | 36 | M | 169 | 79 | 27.7 | 1.9 |
| S2 | 63 | M | 180 | 80 | 24.7 | 2.0 |
| S3 | 34 | M | 175 | 122 | 39.8 | 2.7 |
| S4 | 25 | M | 166 | 56 | 20.3 | 1.6 |
| S5 | 16 | M | 175 | 86 | 28.1 | 2.2 |

To quantify inter-individual anatomical variability within the cohort, chamber volumes derived from the segmented end-diastolic geometries were extracted for all four chambers. Left ventricular (LV), left atrial (LA), right ventricular (RV), and right atrial (RA) volumes are summarised in Table 2.

**Table 2.** End-diastolic chamber volumes (LV, LA, RV, RA) for the five healthy subjects derived from cine MRI segmentation

| ID | LV Volume (cm3) | LA Volume (cm3) | RV Volume (cm3) | RA Volume (cm3) |
| --- | --- | --- | --- | --- |
| S1 | 183.9 | 48.7 | 147.1 | 66.1 |
| S2 | 159.4 | 47.3 | 163.5 | 71.2 |
| S3 | 202.7 | 49.1 | 185.8 | 89.5 |
| S4 | 173.8 | 39.1 | 178.9 | 81.2 |
| S5 | 183.6 | 46.0 | 220.3 | 67.9 |

### Image acquisition

Cine magnetic resonance imaging (MRI) was acquired using a Siemens Prisma 3T scanner (software version VE11C) with a steady-state free precession (SSFP) imaging sequence. Short-axis and long-axis views were obtained to enable full reconstruction of the four cardiac chambers: left ventricle (LV), right ventricle (RV), left atrium (LA), and right atrium (RA), as well as the main vessels. The in-plane spatial resolution was approximately 1.4 mm. Slice thickness differed between chambers, with 4 mm slices used for atrial imaging and 8 mm slices for ventricular imaging. Each cardiac phase consisted of 25 temporal frames. Fig 1 (A1) demonstrates an example of long axis view of one subject.

**Fig 1.**
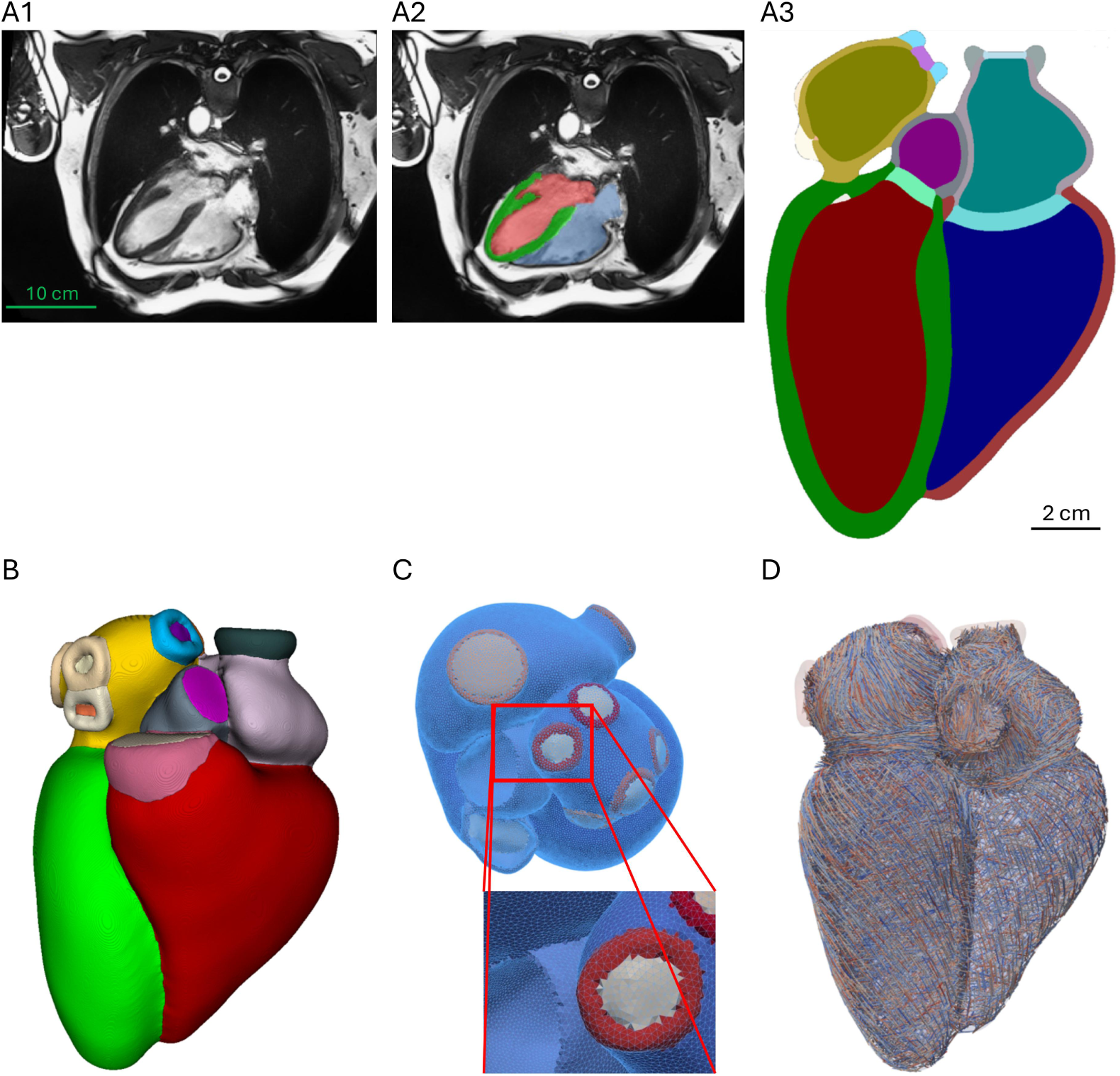
Cine MRI–based four-chamber heart model construction pipeline. (A1) Representative long-axis cine MRI slice at end diastole from one healthy subject. (A2–A3) Anatomical labelling and segmentation of cardiac chambers and major vessels. (B) Three-dimensional reconstruction of the four-chamber heart geometry for one subject. (C) Unstructured tetrahedral mesh of the whole heart with approximately 1 mm mean edge length. (D) Rule-based myocardial fibre assignment in ventricles and atria.

### Segmentation and 3D Reconstruction

Labelling (Fig 1 (A2)) and segmentation (Fig 1 (A3)) of the cardiac structures were performed using a hybrid deep-learning and semi-manual pipeline. A transformer–CNN hybrid network was employed to segment atria, ventricles, and main vessels. The encoder used Swin transformer blocks with cross-window attention [16, 17], while the decoder comprised 3D CNN layers. A Label-Completion U-Net (LC-UNet) was used to reconstruct a complete 3D segmentation from structures incompletely captured in the cine views, enabling recovery of anatomical regions not fully covered by the MRI slices [18]. Additional anatomical completion and post-processing, including generation of thin atrial walls and valve planes, followed the automated four-chamber reconstruction pipeline described in [19]. The locations of the main inlets (superior and inferior vena cava, pulmonary veins) and outlets (aorta, pulmonary artery) were automatically generated by the reconstruction pipeline. The resulting geometries were subsequently inspected and, where necessary, minor adjustments were performed using 3D Slicer [20], and ITK-SNAP [21] to ensure anatomical consistency of inlet and outlet positioning. In total, 31 anatomical labels were reconstructed, including myocardium, blood pools, valves, inflow/outflow tracts, and major vessels. A 3D reconstruction example for one subject is shown in Fig 1B. The reconstructed geometries for all five subjects are shown in S1 Fig.

### Model Creation

This workflow followed the publicly available four-chamber heart cohort pipeline described by [22] and builds upon similar anatomical modelling frameworks in cardiac digital twin studies [13, 23].

### Mesh generation

Segmented four-chamber geometries were meshed using unstructured tetrahedral elements with a target mean edge length of approximately ∼1 mm. Prior to meshing, the geometries were smoothed using Segtools [24]. Tetrahedral meshes were then generated using the CGAL (Computational Geometry Algorithms Library) tetrahedral meshing framework following the four-chamber heart modelling pipeline described in previous studies [19]. The initial segmentations included multiple anatomical labels such as myocardium and blood pools; however, blood pools were excluded from the finite-element mesh used for simulations. Fig 1C shows an example of the meshed whole-heart geometry with a zoomed region illustrating mesh resolution. The meshed geometries of all five subjects are shown in S2 Fig. The resulting finite-element meshed geometries for the five subjects will be made publicly available alongside this work.

### Fibre assignment

Myocardial fibre orientations were defined using a rule-based approach [25, 26]. In the ventricles, fibre angles rotated transmurally from +60 at the endocardium to −60 at the epicardium. Atrial fibres were assigned separately, following longitudinal patterns along the walls and circumferential bundles near the appendages and venous inlets [27]. Fig 1D shows fibre assignment to the myocardium of a subject.

### Mesh Quality Assessment

We assessed tetrahedral element quality via the Scaled Jacobian (SJ), a standard mesh-quality metric that quantifies element distortion by measuring the deviation of the local element mapping from an ideal tetrahedron, with values closer to 1 indicating better element quality and values approaching 0 indicating increasingly distorted elements. Across the five subjects, median SJ ranged from 0.734 to 0.737, with mean ± SD from 0.724 ± 0.111 to 0.727 ± 0.111. The proportion of elements with SJ *<* 0.2 was low (*<* 0.005%), indicating overall good mesh quality suitable for large-deformation mechanics. A summary of mesh quality metrics for all five subjects is provided in Table 3. Per-subject SJ distributions are shown in S3 Fig.

**Table 3.** Mesh quality metrics (Scaled Jacobian) for control hearts

| ID | No. of Elements | Mean | SD | % SJ $< 0.2$ |
| --- | --- | --- | --- | --- |
| S1 | 1,761,995 | 0.726 | 0.111 | 0.002 |
| S2 | 2,018,644 | 0.726 | 0.111 | 0.003 |
| S3 | 2,155,677 | 0.727 | 0.111 | 0.004 |
| S4 | 1,781,396 | 0.724 | 0.111 | 0.003 |
| S5 | 1,941,956 | 0.725 | 0.111 | 0.003 |

### Modelling and simulation

We used a four-chamber electromechanics framework that couples tissue electrophysiology to active tension and large-deformation mechanics, with circulatory and pericardial constraints [13, 28].

Myocardial activation times were computed with a reaction–eikonal formulation [29]. Ventricular and atrial ionic currents were simulated with the ToR-ORd model and the Courtemanche model, respectively; intracellular Ca^2+^ was used to calculate active tension via the Land model in four chambers [13, 30–33]. Fast endocardial conduction was modelled as a thin endocardial layer in the ventricles, extending up to 70% along the apico-basal direction in the ventricles. Bachmann bundle was defined as a fast-conducting inter-atrial region between the LA and RA, generated from the atrial coordinate-based workflow and assigned a higher conduction velocity than the surrounding atrial myocardium [13, 29, 34, 35].

For passive mechanics, atrial and ventricular myocardium were modelled as transversely isotropic hyperelastic tissue using the Guccione strain-energy function. Vessels and valve planes were assigned neo-Hookean material properties. Active stress followed the Land excitation–contraction model and was applied along the locally assigned myocardial fibre direction [13, 32, 36].

The effect of the pericardium was modelled with spatially varying normal springs that penalised outward displacement of the epicardium in the surface-normal direction. The stiffness was scaled so that the apex had the maximum penalty and the base was free, allowing physiological atrioventricular-plane motion. Omni-directional springs were also applied at the pulmonary vein and superior vena cava inlet rings, corresponding to the circular boundaries where these vessels connect to the atrial chambers [22, 23]. The ventricular cavities and great vessels were coupled to a closed-loop 0D circulation model (CircAdapt), which provided preload, afterload, and valve dynamics [13, 28].

### Parameter Selection

The electromechanical heart model comprises a large number of parameters spanning cellular electrophysiology, calcium handling, tissue mechanics, and circulatory boundary conditions. In total, the underlying multiscale framework contains over 100 physiological and numerical parameters, many of which are fixed based on prior studies or physiological measurements. In the present study, we focused on a subset of 46 parameters that are most relevant for global cardiac function.

Following the multiscale sensitivity analysis framework and parameter ranges described in [13], and informed by prior cellular-, passive mechanics-, whole-heart activation time-, and cardiovascular system sensitivity analyses, we selected 46 cardiac input parameters for the present GSA. In addition, we defined 20 output metrics describing global cardiac function, including pressure- and volume-based measures across all four chambers. Sensitivity results for electrophysiological outputs (atrial and ventricular activation times) are summarised in S4 Fig.

To keep the main text focused, we report here the mechanics, CircAdapt, and electrophysiology parameters used in the Latin Hypercube Sampling (LHS), alongside their ranges (Table 4A and Table 4B). The remaining inputs (cellular electrophysiology and excitation–contraction parameters) listed in [13] are provided in S1 Table.

**Table 4A.** Input parameters (Mechanics and CircAdapt) and ranges (Pressures are given in mmHg; valve and orifice areas in mm^2^; Rsys and Rpulm and KArt are expressed in model resistance units consistent with the CircAdapt implementation)

| Parameter | Description | Range (min–max) |
| --- | --- | --- |
| **Mechanics** |  |  |
| a_ventricles | Ventricle stiffness scaling | 0.5 – 5.0 |
| a_atria | Atria stiffness scaling | 1.5 – 6.0 |
| k_peri (MPa/mm) | Pericardial spring stiffness | 0.0005 – 0.05 |
| Tref_lvrv | Reference tension scaling (LV/RV) | 0.5 – 1.0 |
| EDP_lv (mmHg) | LV end-diastolic pressure | 3.0 – 15.0 |
| EDP_rv (mmHg) | RV end-diastolic pressure | 1.6 – 9.6 |
| **CircAdapt** |  |  |
| Rsys | Systemic resistance scaling factor | 0.1 – 5.0 |
| Rpulm | Pulmonary resistance scaling factor | 0.1 – 5.0 |
| mitral_area (mm<sup>2</sup>) | Mitral valve orifice area | 250 – 1500 |
| tricuspid_area (mm<sup>2</sup>) | Tricuspid valve orifice area | 250 – 1500 |
| aortic_area (mm<sup>2</sup>) | Aortic valve orifice area | 150 – 1000 |
| sysOrifice_area (mm<sup>2</sup>) | Systemic outlet area | 150 – 1000 |
| pulmOrifice_area (mm<sup>2</sup>) | Pulmonary outlet area | 150 – 1000 |
| AoPref (mmHg) | Aortic pressure reference | 57 – 160 |
| PaPref (mmHg) | Pulmonary arterial pressure ref. | 4 – 79 |
| VePref (mmHg) | Venous pressure reference | 0.45 – 24 |
| kArt | Arterial compliance/stiffness | 4 – 12 |

**Table 4B.** Electrophysiology input ranges per subject (obtained after history matching)

| Subject | CV_ventricles (m/s) | CV_atria (m/s) | AV_delay (ms) |
| --- | --- | --- | --- |
| S1 | 0.50 – 1.03 | 0.58 – 1.03 | 100 – 200 |
| S2 | 0.69 – 1.03 | 0.54 – 1.03 | 100 – 200 |
| S3 | 0.55 – 1.01 | 0.57 – 1.03 | 100 – 200 |
| S4 | 0.59 – 0.98 | 0.53 – 1.03 | 100 – 200 |
| S5 | 0.69 – 1.03 | 0.54 – 1.03 | 100 – 200 |

Electrophysiological parameter ranges differed per subject (Table 4B) because we first constrained each subject’s electrophysiological parameter space using history matching informed by propensity-matched QRS and P-wave durations (see Supplementary Material: Propensity-Based Calibration of Electrophysiological Parameters). Output parameters are listed in Table 5. Because the electrophysiological parameter ranges were pre-constrained using history matching to propensity-matched ECG targets, the subsequent global sensitivity analysis of pressure- and volume-based outputs should be interpreted as conditional on these physiologically restricted electrical parameter ranges.

**Table 5.**
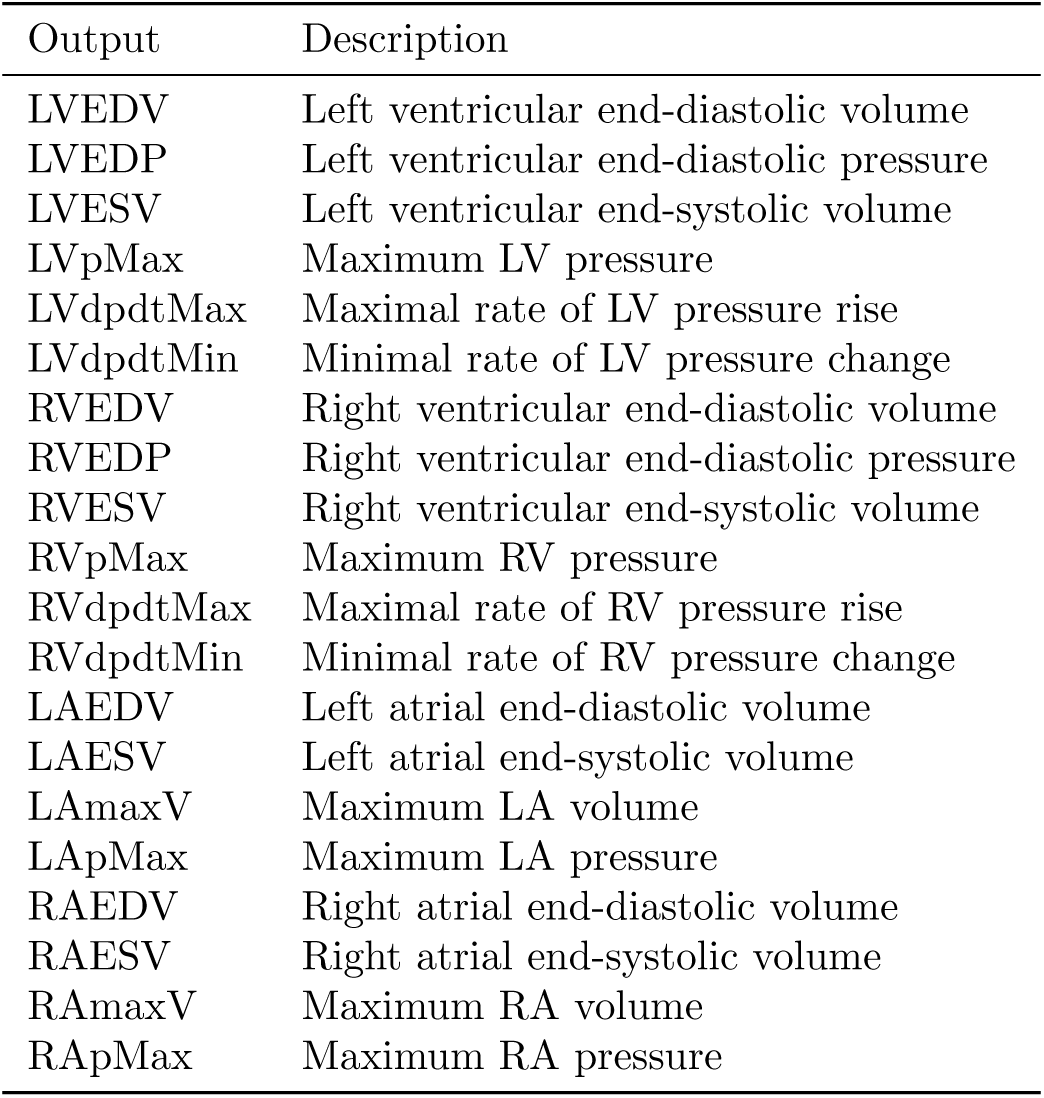
Output parameters.

### Experimental Design

Parameter ranges were sampled using LHS with 500 simulation scenarios per heart. This sampling size is consistent with common recommendations in global sensitivity and emulator studies, which suggest using approximately 10 samples per input parameter to adequately explore the parameter space [37]. With 46 parameters, this corresponds to approximately 460 simulations, and we therefore used 500 samples per subject. For the five healthy subjects, this resulted in a total of 2,500 electromechanical simulations. All simulations were performed on the ARCHER UK National Supercomputing Service using the CARPentry (CARP) software framework [13, 28, 38, 39].

### Emulators, Sensitivity Analysis, and History Matching

GSA of whole-heart electromechanical models requires a large number of simulations across the parameter space. However, each electromechanical simulation is computationally expensive, typically requiring several hours on high-performance computing resources using multiple CPU cores. To make the sensitivity analysis computationally tractable, surrogate models were constructed using GPEs.

GPEs were trained for each output [13, 40]. Training data were generated from LHS of the parameter space. Emulator accuracy was assessed using 5-fold cross-validation, computing the coefficient of determination (*R*^2^) and the independent standard error (ISE) between simulator outputs and emulator predictions. The *R*^2^ metric quantifies pointwise prediction accuracy, while the ISE evaluates the calibration of emulator uncertainty by measuring the percentage of predictions lying within two standard deviations of the emulator mean prediction. The GPEs performance is shown in supplementary S5 Fig for all five subjects.

GSA was performed with Sobol variance-based indices using Saltelli sampling [41]. Total Sobol indices were computed for ranking of the input parameters according to their impact on the 20 outputs. To restrict analysis to physiologically plausible regions, we applied iterative History Matching [40]. Emulator predictions were compared against physiological target ranges, and implausible regions of parameter space were progressively excluded across three successive waves. At each wave, emulator predictions were evaluated against the physiological targets and parameter sets inconsistent with these ranges were discarded. After the final HM wave, GPEs were retrained on the remaining plausible parameter region and used for the final sensitivity analysis. Comparisons were then made across the five hearts to assess the robustness of sensitivity rankings.

### Quantification of Ventricular–Atrial Coupling

To further investigate atrioventricular coupling at the chamber level, we performed a focused post-processing analysis of the global sensitivity results by grouping input parameters according to anatomical specificity. All atrial-specific and all ventricular-specific parameters were included, independent of physiological subsystem (electrophysiological, mechanical, or calcium-handling).

Total Sobol sensitivity indices (*S_T_*) obtained from the GPE-based GSA were used. The analysis was restricted to atrial outputs (LAEDV, LAESV, LAmaxV, LApMax, RAEDV, RAESV, RAmaxV, RApMax).

Input parameters were partitioned into three groups:

1. Ventricular-specific parameters: all parameters associated with ventricular electrophysiology (ToR-ORd model), ventricular mechanics (a ventricles, Tref lvrv), ventricular conduction (CV ventricles), and ventricular filling pressures (EDP lv, EDP rv).
2. Atrial-specific parameters: all parameters associated with atrial electrophysiology (Courtemanche model), atrial passive stiffness (a atria), and atrial conduction (CV atria).
3. ther (global/haemodynamic) parameters: circulatory resistances (Rsys, Rpulm), vascular pressure reference parameters, valve parameters, arterial stiffness, and the pericardial constraint.

For each atrial output, total Sobol indices within each group were summed and renormalised such that the combined contribution of the three groups equalled unity.

## Results

A total of 2,500 electromechanical simulations (500 per subject across five subjects) were performed following LHS of the selected parameter space. GPEs demonstrated high predictive accuracy across all outputs (S5 Fig), enabling efficient GSA within physiologically constrained regions identified through history matching. The cohort consisted of five healthy male subjects spanning a range of ages and body sizes (Table 1). All simulations were performed on high-quality meshes verified using Scaled Jacobian metrics (Table 3 and S3 Fig).

### Pressure–Volume Dynamics

Pressure–volume (PV) loops obtained from the simulations demonstrated stable and physiologically coherent behaviour across all four chambers (S6 Fig). End-diastolic chamber volumes varied across subjects (Table 2), reflecting inter-individual anatomical differences captured in the reconstructed geometries. For each subject, the simulated loops formed well-defined trajectories with chamber-specific morphology characteristic of ventricular filling and ejection phases and atrial reservoir–conduit dynamics. Loop orientation and phase transitions were preserved across the explored parameter space.

To support the qualitative assessment of PV behaviour, key PV-derived biomarkers were quantified for each subject (S2 Table). Across subjects, the median left ventricular end-diastolic volume (LVEDV) ranged from 150 to 205 mL, with within-subject 5th–95th percentile intervals typically spanning approximately 130–250 mL. These values overlap with the upper physiological ranges reported for adult males in large cardiovascular magnetic resonance cohorts [42] and echocardiographic chamber quantification guidelines [43].

Median left ventricular peak pressure (LVpMax) ranged from 99 to 108 mmHg, consistent with normal systemic systolic pressures reported in healthy adults [44, 45]. Right ventricular peak pressure (RVpMax) ranged from 37 to 45 mmHg, preserving the expected lower-pressure pulmonary circulation relative to the systemic circulation. Although slightly above typical resting right ventricular systolic pressures (∼15–30 mmHg), these values remain within physiologically plausible ranges and are consistent with clinically reported thresholds, where right ventricular systolic pressure around or above 40 mmHg is considered mildly elevated [44–46].

Atrial pressures were similarly within physiologically plausible ranges. Median left atrial maximum pressure (LApMax) ranged from 11 to 14 mmHg, consistent with reported left-sided filling pressures in haemodynamic studies [44, 45]. Right atrial maximum pressure (RApMax) ranged from 10 to 11 mmHg across subjects, which is slightly above typical resting right atrial pressures estimated in clinical practice, but remains within physiologically plausible limits, noting that guideline-reported values are based on mean right atrial pressure rather than peak pressure [47].

Left ventricular contractility, assessed via LV dP/dt max, demonstrated median values between 1430 and 1650 mmHg/s. These values fall within the range of left ventricular pressure rise reported in catheter-based and Doppler-validated studies [48].

Although moderate variability in loop width, slope, and enclosed area was observed within and between subjects reflecting differences in compliance, loading conditions, and sampled parameter combinations, the overall magnitudes of pressures and volumes remained within physiologically plausible ranges for healthy adult hearts.

### Global Sensitivity Patterns Across Subjects

Fig 2A shows an example heatmap (Subject S2) illustrating the normalised total effects of 46 input parameters on 20 cardiac outputs. In this heatmap, each row corresponds to a model output, and each column corresponds to an input parameter, while darker colours indicate a stronger influence of that parameter on the given output. Similar sensitivity patterns were observed across all subjects (S7 Fig). The dominant sensitivities consistently clustered around haemodynamic loading parameters. These included systemic and pulmonary resistances (Rsys and Rpulm), left and right ventricular end-diastolic pressures (EDP lv and EDP rv), and the venous reference pressure parameter (VePref), which defines the baseline pressure at the pulmonary venous inflow to the left atrium.

**Fig 2.**
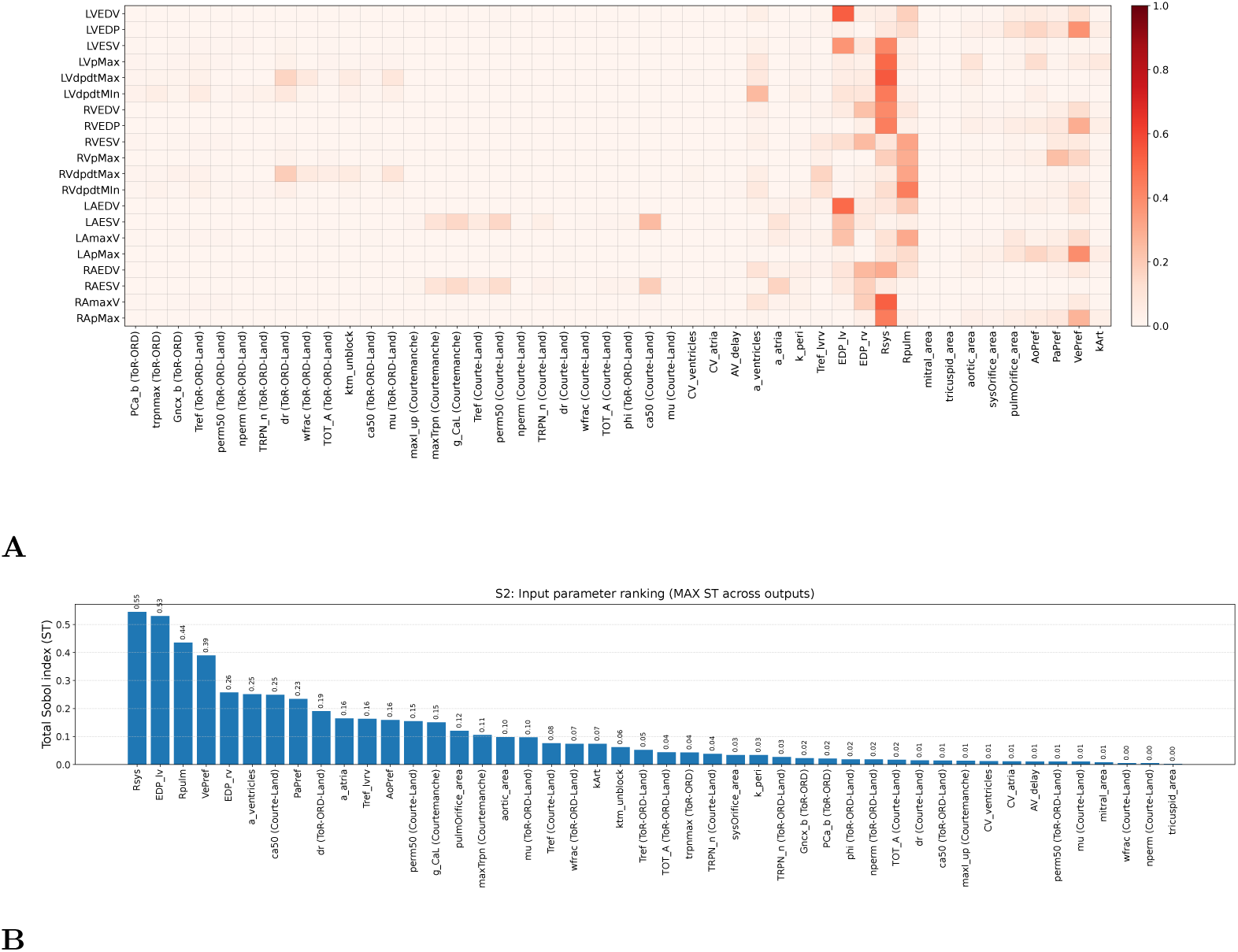
Sensitivity analysis results. (A) Heat map showing the total impact of each input parameter on individual output parameters. (B) Ranking of input parameters based on their maximum total impact.

Parameter ranking plots (Fig 2B) confirmed that these parameters consistently ranked among the top contributors across all subjects. Together, these findings indicate that, in healthy hearts under resting conditions, global cardiac function is primarily governed by preload and afterload conditions rather than by variations in intrinsic tissue properties or electrophysiological parameters. Electrophysiological parameters, such as atrial and ventricular conduction velocities, exhibited comparatively lower sensitivity and predominantly influenced activation timing rather than PV relationships.

### Contributions of Ventricular, Atrial, and Global Parameters to Atrial Outputs

Chamber-level contributions to atrial outputs are summarised in Fig 3. Individual subject results are shown in Fig 3A, while the cohort mean across all five healthy subjects is presented in Fig 3B.

**Fig 3.**
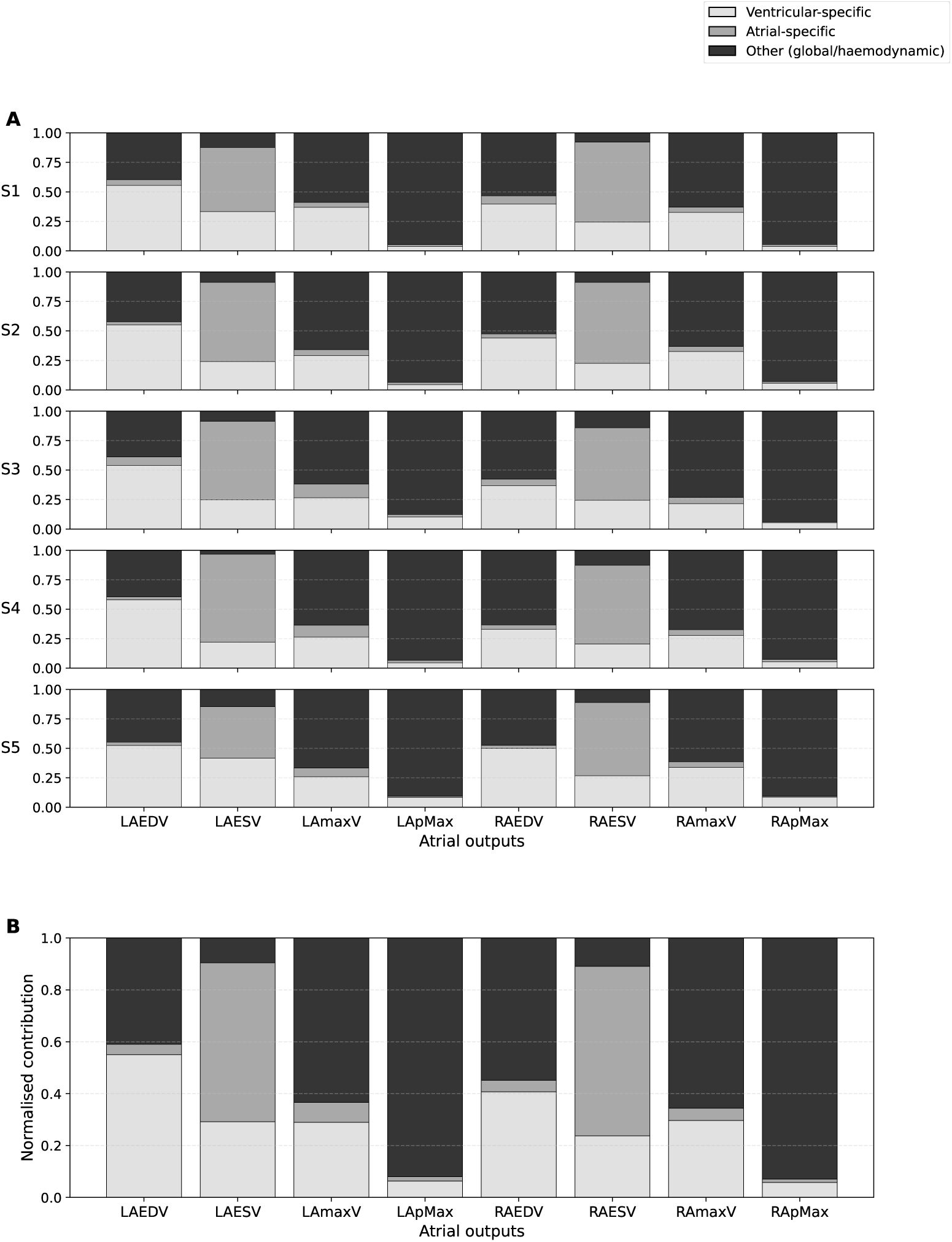
Chamber-level contributions to atrial outputs across five healthy subjects. (A) Normalised total Sobol sensitivity contributions for each atrial output in individual subjects (S1–S5). (B) Cohort mean contributions across all five subjects.

Across all subjects, atrial pressure outputs (LApMax and RApMax) were overwhelmingly dominated by non–chamber-specific haemodynamic parameters, which accounted for more than 90% of total sensitivity on average (LApMax: 92.0 *±* 2.9%; RApMax: 92.9 *±* 1.5%). Ventricular-specific parameters contributed modestly (approximately 5–6%), while atrial-specific parameters had minimal influence (*<* 2%). This indicates that atrial peak pressures in healthy hearts are primarily governed by global loading conditions and vascular boundary parameters rather than intrinsic atrial or ventricular tissue properties.

For atrial end-diastolic volumes (LAEDV and RAEDV), ventricular-specific parameters contributed substantially (LAEDV: 55.0 ± 2.0%; RAEDV: 40.7 ± 6.5%), reflecting the influence of ventricular filling pressures and myocardial properties on atrial preload dynamics. Global haemodynamic parameters also remained important contributors, particularly for the right atrium.

In contrast, atrial end-systolic volumes (LAESV and RAESV) showed the strongest atrial-specific contribution. Atrial-specific parameters accounted for approximately 61.3 ± 12.2% (LAESV) and 65.4 ± 3.4% (RAESV) of total sensitivity, exceeding ventricular-specific contributions (29.2 ± 8.1% and 23.7 ± 2.4%, respectively). This indicates that intrinsic atrial myocardial properties are the primary determinants of atrial emptying, while ventricular and global parameters play secondary roles.

Importantly, the qualitative pattern was consistent across all five subjects (Fig 3A), and the cohort-averaged results (Fig 3B) demonstrate a structured, phase-dependent coupling behaviour: global parameters dominate atrial pressure regulation, ventricular parameters substantially influence atrial filling, and intrinsic atrial properties primarily govern atrial emptying.

## Discussion

### Cohort-Level Parameter Consistency

Across the five healthy subjects, total Sobol index distributions demonstrated a high degree of consistency. Systemic and pulmonary resistances, ventricular end-diastolic pressures, and the venous reference pressure at the left atrium were consistently identified as the most influential parameters of cardiac function. VePref defines the baseline pulmonary venous pressure and therefore directly modulates left atrial filling and downstream ventricular preload. Its importance alongside vascular resistances and ventricular filling pressures highlights the dominant role of haemodynamic boundary conditions in shaping global cardiac function.

The consistency of these sensitivity rankings across anatomically different hearts suggests that organ-scale cardiac behaviour in healthy subjects is primarily controlled by loading conditions rather than by inter-individual differences in geometry or electrophysiological properties. Electrophysiological parameters, including conduction velocities, showed comparatively lower influence on pressure- and volume-based outputs, with their effects mainly affecting timing metrics. This finding reflects the physiological parameter ranges explored in healthy hearts; under pathological conditions or substantial conduction abnormalities, electrophysiological parameters may exert a stronger influence on global haemodynamics.

### Comparison to Previous Studies

These findings are broadly consistent with [13], who identified the above-mentioned parameters as dominant in a single-patient electromechanical model of a female patient with atrial fibrillation and heart failure. Our cohort analysis extends these conclusions by showing that the aforementioned groups of parameters consistently emerge as the most influential across healthy hearts with varying geometry. This indicates that haemodynamic and loading parameters exert the primary control over cardiac function at the organ scale, while electrophysiological variations primarily modulate activation timing.

Our findings are also in agreement with the observations of [27], who examined the link between anatomical variability and simulated cardiac function using statistical shape models. Their study showed that shape modes accounting for a large proportion of anatomical variance do not necessarily explain a comparable proportion of electromechanical functional variability, whereas modes capturing more subtle anatomical features can have a measurable influence on specific functional outputs. These results indicate that the relationship between cardiac anatomy and function is not straightforward, and that variations in anatomy do not uniformly translate into changes in global functional parameters.

Taken together with our cohort-based sensitivity analysis, this suggests that while anatomical structure remains an important feature of cardiac behaviour, its influence on the relative importance of model parameters is limited and that findings of important parameters in a subset of a cohort are potentially generalisable across all patients without the need for patient specific sensitivity analysis.

More broadly, the dominance of loading and boundary-condition parameters in our results is consistent with sensitivity analysis findings across a range of cardiovascular modelling studies. In coupled electromechanical models, variance-based analyses have shown that organ-level biomarkers can be strongly influenced by passive and loading-related parameters when outputs reflect chamber pressures and volumes [12]. Similarly, surrogate-based sensitivity analysis of four-chamber haemodynamic has highlighted substantial control by vascular resistances and circulation parameters on pressure- and flow-related measures [14]. Conversely, electrophysiology-focused sensitivity and uncertainty quantification studies typically identify conduction and activation-related parameters as dominant drivers of ECG and timing metrics, rather than global PV-derived outputs [15]. Taken together, these studies suggest a consistent pattern across modelling frameworks: boundary conditions and filling dynamics dominate pressure/volume biomarkers, whereas electrical parameters primarily govern activation timing, consistent with the separation of sensitivities observed in our cohort.

### Interpretation of Ventricular–Atrial Coupling Mechanisms

These findings suggest a clear functional separation between atrial pressure regulation and atrial volume dynamics. Atrial pressures appear to be predominantly governed by global haemodynamic parameters, particularly vascular resistances and venous reference pressure, highlighting the dominant role of boundary conditions in determining pressure responses. Ventricular-specific parameters contribute modestly, while intrinsic atrial parameters play only a minor role in pressure regulation. This interpretation is consistent with the concept that atrial pressure behaviour reflects the broader haemodynamic environment and the dynamic interaction between atria and ventricles during the cardiac cycle [49].

In contrast, atrial end-systolic volumes are primarily influenced by atrial-specific parameters, indicating that intrinsic atrial myocardial properties are the main determinants of atrial emptying. Ventricular parameters remain relevant but are secondary to atrial intrinsic control in this phase. Atrial end-diastolic volumes reflect an intermediate regime, where ventricular filling pressures and global loading conditions exert substantial influence. This phase-dependent pattern suggests that atrioventricular coupling in healthy hearts is not uniformly ventricular-dominant but instead depends on the functional state of the atrium: global loading governs pressures, ventricular mechanics shape filling, and intrinsic atrial properties dominate emptying. Previous physiological studies have similarly described atrial conduit behaviour as an integrated manifestation of atrioventricular coupling and ventricular filling dynamics [50].

The preservation of this structured coupling pattern across both atria and across all five healthy subjects suggests that coordinated chamber interaction is a defining characteristic of normal cardiac physiology. Alterations in this balance, such as excessive atrial-specific dominance in pressure regulation or diminished ventricular influence on filling, may therefore serve as potential markers of pathological remodelling, particularly in conditions where atrioventricular coupling becomes disrupted [49].

### Limitations

While the present framework provides a systematic evaluation of parameter influence in healthy four-chamber electromechanical models, several considerations should be noted when interpreting the findings.

First, the cohort consisted exclusively of male healthy control subjects. Although the five anatomies span a broad range of ages, body mass indices, and chamber volumes, sex-specific physiological differences were not explicitly evaluated. However, the range of cardiac sizes represented in this cohort overlaps with the physiological spectrum reported for both male and female hearts in large imaging cohorts such as the UK Biobank. Moreover, the dominant parameter groups identified here are consistent with previous electromechanical sensitivity analyses performed in a heart-failure model of a female patient with atrial fibrillation [13]. While the present study did not explicitly evaluate female subjects, the consistency of sensitivity rankings across hearts of different anatomical sizes and across healthy and pathological models suggests that these findings are unlikely to be strictly sex-specific.

Second, all models represent healthy hearts. This provides a controlled baseline for evaluating parameter influence under physiological conditions. However, sensitivity patterns may differ in pathological states where myocardial stiffness, contractility, or loading conditions are altered beyond the physiological ranges evaluated here. The dominance of haemodynamic loading parameters observed here should therefore be interpreted primarily within the context of physiological parameter ranges. Nevertheless, the consistency of dominant parameters with those identified in the previously studied atrial-fibrillation heart-failure case [13] suggests that similar sensitivity structures may extend to certain pathological settings.

Third, the cardiovascular model does not incorporate dynamic regulatory mechanisms such as baroreflex feedback, autonomic nervous system modulation, or sympathetic activation. In vivo, these regulatory pathways continuously adjust heart rate, contractility, and vascular tone in response to haemodynamic perturbations. The absence of such feedback mechanisms means that the present simulations reflect a mechanically and electrically coupled system without neurohumoral physiological control. Although the closed-loop CircAdapt circulation model dynamically updates haemodynamic boundary conditions, the overall system does not include adaptive autonomic regulation. Inclusion of such physiological control mechanisms could modify sensitivity rankings, particularly for pressure-related outputs.

Fourth, parts of the anatomical reconstruction relied on a label-completion network to infer structures not fully captured in the cine MRI views. Because these regions are not directly observed in the imaging data, the reconstruction process may introduce geometric uncertainty in these structures.

Finally, electrophysiological parameter ranges were constrained through history matching to propensity-matched ECG features derived from the UK Biobank cohort rather than fully subject-specific electrical mapping. In particular, electrophysiological parameter ranges were restricted to reproduce physiologically plausible QRS and P-wave durations observed in propensity-matched subjects. Although this approach ensured realistic activation dynamics and reduced implausible regions of parameter space, it may reduce the apparent influence of electrophysiological parameters on PV outputs.

Despite these considerations, the cohort encompasses a substantial range of anatomical sizes and loading conditions and shows sensitivity patterns that are broadly consistent with those previously reported in a female atrial-fibrillation heart-failure model [13], supporting the robustness of the observed parameter rankings across different heart sizes, disease states, and subject characteristics.

## Conclusions

We have presented a repeatable cine MRI–based workflow for constructing four-chamber electromechanical heart models and applied cohort-level GSA in five healthy subjects. Across anatomically distinct hearts, within the physiologically restricted parameter space defined after history matching, haemodynamic loading parameters, including systemic and pulmonary resistances, ventricular filling pressures, and venous inflow pressure, consistently emerged as dominant determinants of pressure- and volume-based cardiac outputs. This highlights the central role of boundary conditions and preload–afterload balance in governing organ-scale cardiac function under healthy resting conditions. The consistency of sensitivity rankings across subjects suggests that, within this small cohort of healthy male subjects spanning a range of body sizes and chamber volumes, the relative importance of major model parameters is relatively robust to the anatomical variability represented here.

A chamber-level ventricular–atrial coupling analysis further revealed a structured, phase-dependent interaction between atria and ventricles. Atrial pressures were primarily governed by global haemodynamic parameters, atrial filling volumes were strongly influenced by ventricular-specific properties, and atrial emptying showed greater dependence on intrinsic atrial parameters. This separation of pressure regulation, filling dynamics, and emptying behaviour provides a mechanistic interpretation of atrioventricular coupling in healthy hearts.

By providing an openly available set of anatomically detailed meshes with a clearly defined modelling and sensitivity analysis framework, this study establishes a benchmark reference for whole-heart electromechanical modelling in healthy subjects. Identifying the parameters that most strongly influence cardiac performance can help guide model calibration strategies and support the development of more reliable cardiac digital twin frameworks.

## Acknowledgments

This work was supported by the Medical Research Futures Fund (grant number 2017687) and a Cardiovascular Early-Mid Career Researcher Grant from NSW Health. Computational resources were provided by the ARCHER2 UK National Supercomputing Service (https://www.archer2.ac.uk). The authors acknowledge the Imaging Centre at Victor Chang Cardiac Research Institute (VCCRI) and the Advanced Cardiac Imaging Centre at St Vincent’s Hospital for their support with data acquisition. C.R. receives funding from the British Heart Foundation (RG/20/4/34803).

## Data Availability

The finite-element meshed heart geometries for the five subjects analysed in this study are available at Zenodo (https://zenodo.org/records/19203952). The Zenodo record is publicly accessible; however, access to the files is restricted. Requests for access can be made via.

## Supporting information

**S1 Fig.** Segmentation and 3D reconstruction results of the five control hearts. The completed segmentation consisted of 31 anatomical regions, including chamber blood pools, myocardium, major vessel inlets and outlets. Valve planes were represented as surface interfaces separating adjacent chamber cavities. Additional capping surfaces were introduced at the vessel inlets/outlets to ensure the endocardial boundaries were fully closed.

**S2 Fig.** Meshed geometries of the five subjects. Unstructured tetrahedral finite-element meshes generated from the segmented anatomies, with a target mean edge length of approximately 1 mm.

**S3 Fig.** Mesh quality histograms for subjects S1–S5. Distributions of the number of elements as a function of the Scaled Jacobian (SJ) metric. Median SJ ranged from 0.734 to 0.737 across subjects, with less than 0.005% of elements below SJ = 0.2.

**S4 Fig.** Polar plots showing total-order Sobol sensitivity indices for total atrial and ventricular activation times across five subjects. CVf: conduction velocity in the fibre direction; KFEC: scaling factor for the conduction velocity of the fast endocardium (FEC); KBB: scaling factor for the conduction velocity of Bachmann bundle; KAni: anisotropy ratio.

**S5 Fig.** GPE performance across five subjects. Emulator predictions (red points) vs. simulator outputs (black curves) for all 20 cardiac outputs, assessed by 5-fold cross-validation. Panels show subjects S1–S5.

**S6 Fig.** Last-beat pressure–volume loops for the four cardiac chambers across all five subjects. Loops extracted from the final beat of a five-beat simulation.

**S7 Fig.** GSA heatmaps for all subjects. Effect of changes in 46 input parameters on the 20 cardiac output parameters for subjects S1–S5. Darker colours indicate stronger influence.

**S1 Table.** Cell dynamics input parameters. Parameters from the atrial Courtemanche model, the ventricular ToR-ORd model, and the Land active tension model.

**S2 Table.** Median [5–95% range] of simulated PV biomarkers across the five subjects. Values reported across the sampled parameter space after history matching.

**S1 Appendix.** Supplementary methods. Propensity-based calibration of electrophysiological parameters; history matching of electrophysiological parameters; global sensitivity analysis of electrophysiological activation times; Gaussian process emulator performance.

## References

1. Fumagalli I, Pagani S, Vergara C, Adebo DA, Del Greco M, Frontera A, et al. The role of computational methods in cardiovascular medicine: a narrative review. Translational Pediatrics. 2024;13(1):146.

2. Niederer SA, Lumens J, Trayanova NA. Computational models in cardiology. Nature reviews cardiology. 2019;16(2):100–111.

3. Prakosa A, Arevalo HJ, Deng D, Boyle PM, Nikolov PP, Ashikaga H, et al. Personalized virtual-heart technology for guiding the ablation of infarct-related ventricular tachycardia. Nature biomedical engineering. 2018;2(10):732–740.

4. Viola F, Del Corso G, De Paulis R, Verzicco R. GPU accelerated digital twins of the human heart open new routes for cardiovascular research. Scientific reports. 2023;13(1):8230.

5. Salvador M, Strocchi M, Regazzoni F, Augustin CM, Dede’ L, Niederer SA, et al. Whole-heart electromechanical simulations using latent neural ordinary differential equations. NPJ Digital Medicine. 2024;7(1):90.

6. Kong F. Automated Model Construction for Image-Based Cardiac Computational Simulations. University of California, Berkeley; 2022.

7. Sung E, Kyranakis S, Daimee UA, Engels M, Prakosa A, Zhou S, et al. Evaluation of a deep learning-enabled automated computational heart modelling workflow for personalized assessment of ventricular arrhythmias. The Journal of Physiology. 2024;602(18):4625–4644.

8. Sel K, Osman D, Zare F, Masoumi Shahrbabak S, Brattain L, Hahn J, et al. Building digital twins for cardiovascular health: From principles to clinical impact. Journal of the American Heart Association. 2024;13(19):e031981.

9. Niederer SA, Aboelkassem Y, Cantwell CD, Corrado C, Coveney S, Cherry EM, et al. Creation and application of virtual patient cohorts of heart models. Philosophical Transactions of the Royal Society A. 2020;378(2173):20190558.

10. Waight MC, Prakosa A, Li AC, Truong A, Bunce N, Marciniak A, et al. Heart Digital Twins Predict Features of Invasive Reentrant Circuits and Ablation Lesions in Scar-Dependent Ventricular Tachycardia. Circulation: Arrhythmia and Electrophysiology. 2025;18(8):e013660.

11. Ugurlu D, Qian S, Fairweather E, Mauger C, Ruijsink B, Toso LD, et al. Cardiac digital twins at Scale from MRI: open tools and representative models from∼ 55000 UK Biobank participants. Plos one. 2025;20(7):e0327158.

12. Levrero-Florencio F, Margara F, Zacur E, Bueno-Orovio A, Wang ZJ, Santiago A, et al. Sensitivity analysis of a strongly-coupled human-based electromechanical cardiac model: Effect of mechanical parameters on physiologically relevant biomarkers. Computer methods in applied mechanics and engineering. 2020;361:112762.

13. Strocchi M, Longobardi S, Augustin CM, Gsell MAF, Petras A, Rinaldi CA, et al. Cell to whole organ global sensitivity analysis on a four-chamber heart electromechanics model using Gaussian processes emulators. PLOS Computational Biology. 2023;19(6):e1011257.

14. Karabelas E, Longobardi S, Fuchsberger J, Razeghi O, Rodero C, Strocchi M, et al. Global sensitivity analysis of four chamber heart hemodynamics using surrogate models. IEEE Transactions on Biomedical Engineering. 2022;69(10):3216–3223.

15. Winkler B, Nagel C, Farchmin N, Heidenreich S, Loewe A, Dössel O, et al. Global sensitivity analysis and uncertainty quantification for simulated atrial electrocardiograms. Metrology. 2022;3(1):1–28.

16. Qayyum A, Razzak I, Mazher M, Lu X, Niederer SA. Unsupervised unpaired multiple fusion adaptation aided with self-attention generative adversarial network for scar tissues segmentation framework. Information Fusion. 2024;106:102226.

17. Mazher M, Razzak I, Qayyum A, Tanveer M, Beier S, Khan T, et al. Self-supervised spatial–temporal transformer fusion based federated framework for 4D cardiovascular image segmentation. Information Fusion. 2024;106:102256.

18. Xu H, Muffoletto M, Niederer SA, Williams SE, Williams MC, Young AA. Whole heart 3D shape reconstruction from sparse views: leveraging cardiac computed tomography for cardiovascular magnetic resonance. In: International Conference on Functional Imaging and Modeling of the Heart. Springer; 2023. p. 255–264.

19. Solis-Lemus JA, Barrows RK, Rodero C, Strocchi M, Montarello N, Lahoti N, et al. Assessing the Importance of Sex and Disease-Specific Anatomy in Electrophysiology and Mechanical Simulations in a Public Virtual Cohort of Four-Chamber Heart Models. Under review. 2026;.

20. Fedorov A, Beichel R, Kalpathy-Cramer J, Finet J, Fillion-Robin JC, Pujol S, et al. 3D Slicer as an Image Computing Platform for the Quantitative Imaging Network. Magnetic Resonance Imaging. 2012;30(9):1323–1341. Available from: https://www.slicer.org/.

21. Yushkevich PA, Piven J, Cody Hazlett H, Gimpel Smith R, Ho S, Gee JC, et al. User-guided 3D active contour segmentation of anatomical structures: Significantly improved efficiency and reliability. NeuroImage. 2006;31(3):1116–1128. Available from: https://www.itksnap.org/.

22. Strocchi M, Augustin CM, Gsell MAF, Karabelas E, Neic A, Gillette K, et al. A publicly available virtual cohort of four-chamber heart meshes for cardiac electro-mechanics simulations. PloS one. 2020;15(6):e0235145.

23. Strocchi M, Gsell MAF, Augustin CM, Razeghi O, Roney CH, Prassl AJ, et al. Simulating ventricular systolic motion in a four-chamber heart model with spatially varying robin boundary conditions to model the effect of the pericardium. Journal of biomechanics. 2020;101:109645.

24. Buske OJ, Hoffman MM, Ponts N, Le Roch KG, Noble WS. Exploratory analysis of genomic segmentations with Segtools. BMC Bioinformatics. 2011;12:415.

25. Streeter Jr DD, Spotnitz HM, Patel DP, Ross Jr J, Sonnenblick EH. Fiber orientation in the canine left ventricle during diastole and systole. Circulation research. 1969;24(3):339–347.

26. Bayer JD, Blake RC, Plank G, Trayanova NA. A novel rule-based algorithm for assigning myocardial fiber orientation to computational heart models. Annals of biomedical engineering. 2012;40:2243–2254.

27. Rodero C, Strocchi M, Marciniak M, Longobardi S, Whitaker J, O’Neill MD, et al. Linking statistical shape models and simulated function in the healthy adult human heart. PLoS computational biology. 2021;17(4):e1008851.

28. Augustin CM, Gsell MAF, Karabelas E, Willemen E, Prinzen FW, Lumens J, et al. A computationally efficient physiologically comprehensive 3D–0D closed-loop model of the heart and circulation. Computer methods in applied mechanics and engineering. 2021;386:114092.

29. Neic A, Campos FO, Prassl AJ, Niederer SA, Bishop MJ, Vigmond EJ, et al. Efficient computation of electrograms and ECGs in human whole heart simulations using a reaction-eikonal model. Journal of computational physics. 2017;346:191–211.

30. Tomek J, Bueno-Orovio A, Passini E, Zhou X, Minchole A, Britton O, et al. Development, calibration, and validation of a novel human ventricular myocyte model in health, disease, and drug block. elife. 2019;8:e48890.

31. Courtemanche M, Ramirez RJ, Nattel S. Ionic mechanisms underlying human atrial action potential properties: insights from a mathematical model. American Journal of Physiology-Heart and Circulatory Physiology. 1998;275(1):H301–H321.

32. Land S, Park-Holohan SJ, Smith NP, Dos Remedios CG, Kentish JC, Niederer SA. A model of cardiac contraction based on novel measurements of tension development in human cardiomyocytes. Journal of molecular and cellular cardiology. 2017;106:68–83.

33. O’Hara T, Virág L, Varró A, Rudy Y. Simulation of the undiseased human cardiac ventricular action potential: model formulation and experimental validation. PLoS computational biology. 2011;7(5):e1002061.

34. Strocchi M, Lee AWC, Neic A, Bouyssier J, Gillette K, Plank G, et al. His-bundle and left bundle pacing with optimized atrioventricular delay achieve superior electrical synchrony over endocardial and epicardial pacing in left bundle branch block patients. Heart rhythm. 2020;17(11):1922–1929.

35. Lee AWC, Nguyen UC, Razeghi O, Gould J, Sidhu BS, Sieniewicz B, et al. A rule-based method for predicting the electrical activation of the heart with cardiac resynchronization therapy from non-invasive clinical data. Medical image analysis. 2019;57:197–213.

36. Guccione JM, McCulloch AD, Waldman LK. Passive material properties of intact ventricular myocardium determined from a cylindrical model. J Biomech Eng. 1991;113(1):42–55.

37. Loeppky JL, Sacks J, Welch WJ. Choosing the sample size of a computer experiment: A practical guide. Technometrics. 2009;51(4):366–376.

38. Rahmani S, Pouliopoulos J, Lee AWC, Barrows RK, Strocchi M, Rodero C, et al. Cardiac Sensitivities to Biomechanical Changes in a Chronic Alcoholic Heart: A Case Study Using 3-Dimensional Electro-Mechanical Heart Modelling. Computing in Cardiology. 2024;51:1.

39. Vigmond EJ, Aguel F, Trayanova NA. Computational techniques for solving the bidomain equations in three dimensions. IEEE Transactions on Biomedical Engineering. 2002;49(11):1260–1269.

40. Longobardi S, Lewalle A, Coveney S, Sjaastad I, Espe EKS, Louch WE, et al. Predicting left ventricular contractile function via Gaussian process emulation in aortic-banded rats. Philosophical Transactions of the Royal Society A. 2020;378(2173):20190334.

41. Saltelli A, Annoni P, Azzini I, Campolongo F, Ratto M, Tarantola S. Variance based sensitivity analysis of model output. Design and estimator for the total sensitivity index. Computer physics communications. 2010;181(2):259–270.

42. Petersen SE, Aung N, Sanghvi MM, Zemrak F, Fung K, Paiva JM, et al. Reference ranges for cardiac structure and function using cardiovascular magnetic resonance (CMR) in Caucasians from the UK Biobank population cohort. Journal of cardiovascular magnetic resonance. 2016;19(1):18.

43. Lang RM, Badano LP, Mor-Avi V, Afilalo J, Armstrong A, Ernande L, et al. Recommendations for cardiac chamber quantification by echocardiography in adults: an update from the American Society of Echocardiography and the European Association of Cardiovascular Imaging. European Heart Journal-Cardiovascular Imaging. 2015;16(3):233–271.

44. Moscucci M. Grossman & Baim’s cardiac catheterization, angiography, and intervention. Lippincott Williams & Wilkins; 2020.

45. Hall JE, Hall ME. Guyton and Hall textbook of medical physiology e-book: Guyton and Hall textbook of medical physiology e-book. Elsevier Health Sciences; 2020.

46. Kotrri G, Youngson E, Fine NM, Howlett JG, Lyons K, Paterson DI, et al. Right ventricular systolic pressure trajectory as a predictor of hospitalization and mortality in patients with chronic heart failure. CJC open. 2023;5(9):671–679.

47. Rudski LG, Lai WW, Afilalo J, Hua L, Handschumacher MD, Chandrasekaran K, et al. Guidelines for the echocardiographic assessment of the right heart in adults: a report from the American Society of Echocardiography: endorsed by the European Association of Echocardiography, a registered branch of the European Society of Cardiology, and . Journal of the American society of echocardiography. 2010;23(7):685–713.

48. Bargiggia GS, Bertucci C, Recusani F, Raisaro A, De Servi S, Valdes-Cruz LM, et al. A new method for estimating left ventricular dP/dt by continuous wave Doppler-echocardiography. Validation studies at cardiac catheterization. Circulation. 1989;80(5):1287–1292.

49. Di Terlizzi V, Barone R, Di Nunno N, Alcidi G, Brunetti ND, Iacoviello M. The atrioventricular coupling in heart failure: pathophysiological and therapeutic aspects. Reviews in Cardiovascular Medicine. 2024;25(5):169.

50. Marino PN. Left atrial conduit function: A short review. Physiological Reports. 2021;9(19):e15053.

